# Modelling cigarette smoke-induced lung vascular dysfunction using an alveolus-on-chip

**DOI:** 10.1101/2025.10.30.685580

**Authors:** Abilash Ravi, Tarek Gensheimer, Xinhui Wu, Jill R. Johnson, Martin C. Harmsen, Reinoud Gosens, Pieter S. Hiemstra, Andries D. van der Meer, Anne M. van der Does

**Author notes:** Corresponding author Anne M. van der Does, PhD, PulmoScience Lab, Department of Pulmonology, Leiden University Medical Center, P.O. Box 9600, 2300 RC Leiden, the Netherlands. Shared first authors (Abilash Ravi, Tarek Gensheimer). Shared last authors (Andries D. van der Meer, Anne M. van der Does).

## Abstract

The alveolus is central to gas exchange in the lung, and alveolar damage is a characteristics of a variety of lung diseases. Understanding alveolar microvascular dynamics and epithelial-endothelial interactions is essential for accurately modeling alveolar physiology and its dysfunction in lung diseases such as Chronic Obstructive Pulmonary Disease (COPD), pulmonary fibrosis and acute respiratory distress syndrome (ARDS). In this study, we present an open-top, membrane-free alveolus-on-chip platform incorporating self-assembled, perfusable 3D vascular networks by primary human lung endothelial cells and pericytes, co-cultured with alveolar epithelial type 2 (AEC2) cells. These vascular networks were developed within 6 days under continuous flow and remained stable for at least 12 days. The inclusion of pericytes supported capillary-like vessel formation and increased gene expression of *EDNRB1*, a gene enriched in alveolar microvascular endothelial cells. Furthermore, *CD31* gene expression was higher in 3D endothelial networks compared to 2D endothelial monolayers, suggesting increased cell-cell adhesion. Monocytes could be successfully perfused through the networks, expanding the platform’s potential for studying immune interactions in lung disorders. Culturing AEC2 monolayers directly on the vascularized hydrogel enabled physiologically relevant cell-cell interactions without artificial membranes, while maintaining air-liquid interface conditions. Importantly, exposure of the AEC2 layer to whole cigarette smoke (WCS) led to complete disintegration of the underlying vascular network, an effect not observed in the absence of AEC2. This chip model provides a human-relevant system for investigating vascular-epithelial crosstalk in the alveolus, smoke-induced lung injury, and immune recruitment, offering a valuable platform for future disease modeling and drug testing applications.

## Introduction

The alveolar-capillary compartment of the lung represents a dynamic and complex microenvironment dedicated to gas exchange. Cigarette smoke (CS) is the primary risk factor for Chronic Obstructive Pulmonary Disease (COPD), but also other forms of air pollution contribute to its development. One of the consequences of prolonged exposure to CS is progressive alveolar wall deterioration, also known as emphysema [1]. Besides causing direct or indirect damage to the alveolar epithelium, CS exposure is known to disrupt the integrity of the lung microvascular barrier [2], evidenced by a reduced expression of endothelial cell markers in lung tissue from patients with COPD [3].

We recently demonstrated a loss of microvascular endothelial cells in morphologically intact alveolar septa in patients with emphysema. Interestingly, in these tissue areas the percentage of alveolar type 2 epithelial cells (AEC2), responsible for repopulating the epithelium after damage, was not different from controls [3]. These *in vivo* data suggest that the loss of endothelial cells may precede that of the alveolar epithelial cells and that CS exposure may lead to impaired repair of damaged epithelium by negatively affecting the vascular compartment [4]. This highlights the need for more insight into cellular interactions within the alveolar niche during CS exposure.

Besides their role in gas exchange, lung microvascular endothelial cells are hypothesized to contribute to alveolar repair and regeneration. Various studies have demonstrated their contribution to these processes through e.g. secretion of mediators like BMP-6 [5], thrombospondin-1 [6], matrix metalloproteinase-14 and by direct contact-based interaction with the alveolar epithelium [7]. In line, we showed that lung endothelial cells, isolated and expanded *in vitro* from donor-derived lung tissue, support AEC2 organoid formation and that this support function was impaired by cigarette smoke exposure of endothelial cells [3]. Despite a clear effect of CS exposure on the microvasculature of the alveolar compartment [8-10], it is currently unclear if CS exerts these effects directly or indirectly, and which cellular cross-talks are involved in the damaging effects on the microvasculature.

Advanced human cell culture models are increasingly used to study processes related to tissue damage and repair [11]. Several micro-engineered lung-on-chip models have been developed to more closely mimic human lung tissue biology than conventional cell culture and animal models, aiming to improve human disease modelling and thereby serve as an alternative to animal testing [12, 13]. However, it remains challenging to develop and apply human-relevant tissue models that represent the complex environment of the alveolar units in the lower respiratory tract, including controlled gas exposure. Several lung-on-chip models have recapitulated the alveolar niche with a range of physiological complexities, including mechanical stretch to mimic breathing [13] and applying dynamic flow to represent blood perfusion [14]. However, these models rely on endothelial monolayers that do not accurately represent the phenotype, organization or function of pulmonary capillaries [15]. Additionally, there is a lack of models that employ a combination of primary cell-types derived from patients with lung disorders.

To address this gap, we aimed to develop an alveolus-on-chip that incorporates primary lung endothelial cells and pericytes, forming perfusable, three-dimensional, stable networks, in co-culture with primary alveolar epithelial cells. In this study, we tested the feasibility of using human-derived primary endothelial cells from lung tissue to form vascular networks with pericytes (also patient-derived). Next, we assessed the compatibility of co-culturing a monolayer of AEC2 with self-assembled vascular networks to study their interactions in our chip system. By seeding the AEC2 monolayer on top of the endothelial networks in hydrogel, we aimed to develop a membrane-free chip system for direct cell interaction studies. In addition, the open-top design of the cell chamber allowed us to study the effects of exposure to cigarette smoke. Overall, we developed a human-relevant model to mimic cigarette smoke-induced damage in the alveoli, facilitating controllable studies on the role of the lung microvasculature in alveolar repair and regeneration.

## Materials and methods

### Isolation, maintenance and expansion of endothelial cells

Endothelial cells and pericytes were isolated from macroscopically normal lung tissue obtained from patients undergoing resection surgery for lung cancer at the Leiden University Medical Center, the Netherlands. Patients from whom this lung tissue was derived were enrolled in the biobank via a no-objection system for coded anonymous further use of such tissue (www.coreon.org). Samples from this Biobank were approved for research use by the institutional medical ethical committee (BB22.006/AB/ab). Since 01-09-2022, patients are enrolled in the biobank using written informed consent in accordance with local regulations from the LUMC biobank with approval by the institutional medical ethical committee (B20.042/KB/kb). The characteristics of donors involved in the study are summarized in **Table 1**.

**Table 1.**
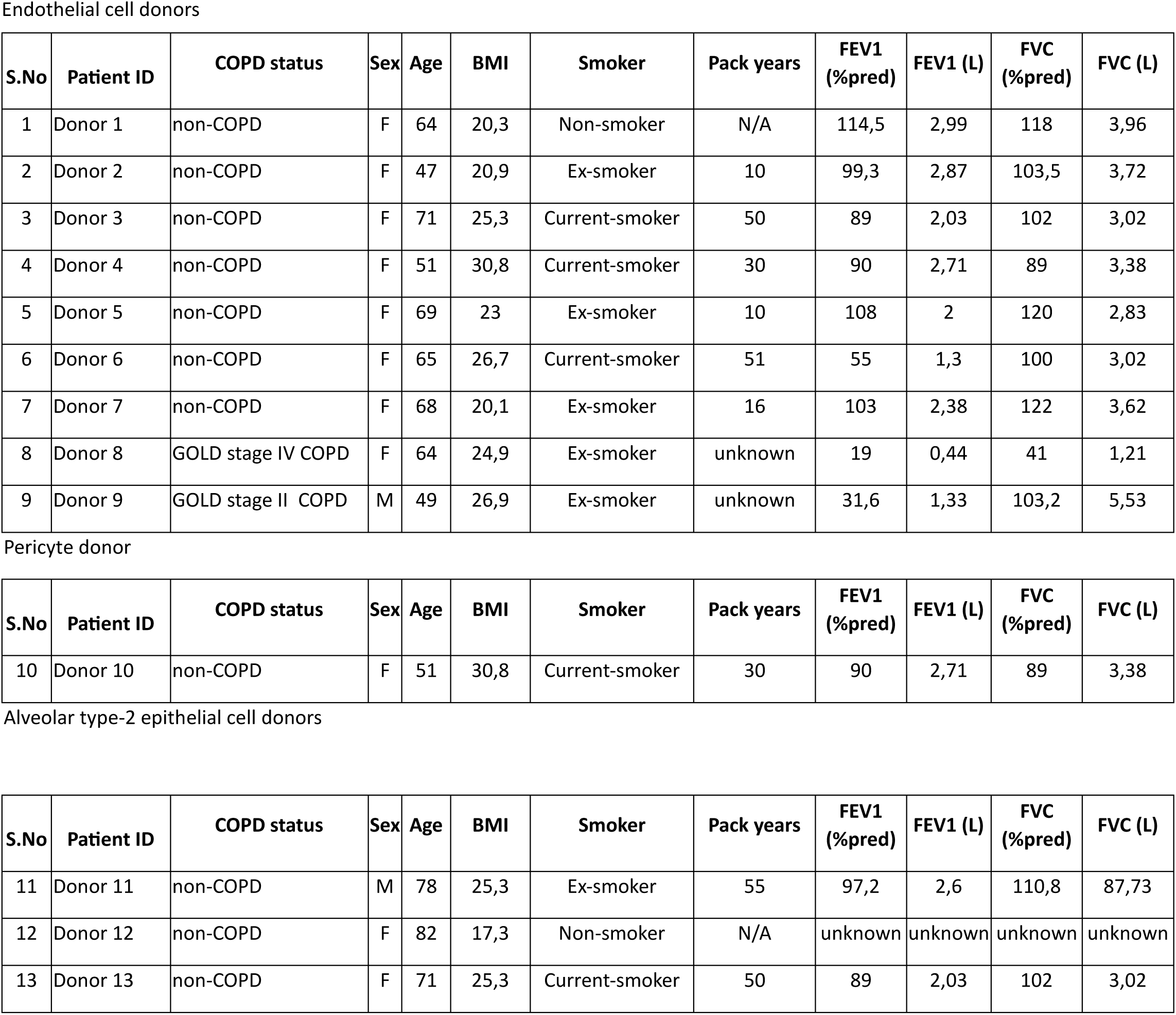
Baseline characteristics of COPD and non-COPD donors Endothelial cell donors.

After rinsing the tissue with Hank’s Balanced Salt Solution (HBSS, Thermo Fisher Scientific, USA), it was cut into small pieces of < 1 mm^3^ and collected in a sterile container. Next, the lung tissue homogenate was digested in 9 ml of 0.1 % collagenase (#07418, STEMCELL Technologies, Canada), 1 ml of 5 U/ml of dispase (#07913, STEMCELL Technologies, Canada) solution and 75 μl of DNase (#DN25-10MG, final concentration: 1 mg/ml; Sigma, USA) per gram of tissue for 1 hour (h) at 37°C. The digested tissue was then collected in gentleMACS™ C tubes (Miltenyi Biotec, the Netherlands) and dissociated twice in the gentleMACS™ Dissociator using program M_lung_2.01 (Miltenyi Biotec, the Netherlands). The homogenate was next passed through a metal sieve, followed by passing through a strainer with a mesh size of 100 μm (VWR International, USA) to obtain a more single cell suspension. The cells were then centrifuged at 265 *g* for 5 min at room temperature (RT) and the pellet was resuspended in 100 μl of EGM-2MV Lonza endothelial medium (#CC-3202, Lonza, Switzerland). To the cell suspension, 40 μl of CD31 microbeads were added (#130-091-935, Miltenyi Biotec, the Netherlands) and endothelial cells were subsequently magnetically sorted according to the manufacturer’s instruction (Miltenyi Biotec, the Netherlands). The endothelial cells were plated in 50X Matrigel-coated (#354230, Corning, USA) plates with the addition of 10 μM SB431542, a TGF-β signalling inhibitor (#S4317, Sigma-Aldrich, USA) and 10 μM Y-27632, a Rho-kinase inhibitor (#10005583, Sanbio B.V., the Netherlands). The endothelial cells received a medium change with endothelial cell culture medium supplemented with 10 μM SB431542, for every two days until they reached confluency.

The expanded endothelial cells were again MACS sorted using CD31 microbeads, after detaching cells in TrypLE^TM^ Select (#12563011, Thermo Fisher Scientific, USA) for 5 min at 37°C. Cells were plated in Matrigel coated-6W plate with endothelial medium supplemented with 10 μM SB431542. For cryopreservation, the confluent monolayer of CD31^+^ endothelial cells were detached using TrypLE^TM^ Select for 5 min at 37°C. After washing the cells in cell culture medium once, cells were resuspended in CryoStor® (#100-1061, STEMCELL Technologies, Canada) and stored in liquid nitrogen. Upon subsequent thawing for chip cultures, endothelial cells were expanded by addition of TGF-β signaling inhibitor SB431542, but was not used for on-chip cultivation.

### Isolation, maintenance and expansion of pericytes

The negative fraction of CD31-sorted cells from the single cell suspension of lung tissue, was used to isolate pericytes. The CD31^-^ fraction was centrifuged at 265 *g* for 5 min at RT. The pellet was resuspended in 100 μl endothelial medium, combined with 2 µl of anti-NG2 primary rabbit IgG antibody (#ab255811, Abcam, USA) and incubated for 15 min at 4°C. Next, the cell suspension was centrifuged, and the pellet was resuspended in 100 μl of endothelial medium with 40 µl of anti-rabbit IgG microbeads (#130-048-602, Miltenyi Biotec, the Netherlands) for 15 min at 4°C. The manufacturer’s instructions were followed for NG2^+^ magnetic selection and the sorted pericytes were plated in endothelial cell culture medium, without SB431542. Cell culture medium was refreshed every two days until confluency. The expanded pericytes were detached using TrypLE^TM^ Select for 5 min at 37°C. Pericytes were washed in cell culture medium, resuspended in CryoStor® and stored in liquid nitrogen until further use.

### Flow cytometry

The expanded endothelial cells and pericytes were characterized using specific markers before seeding in the chip. After thawing and subsequent expansion, the expanded cells were detached using TrypLE^TM^ Select for 5 min at 37°C and washed once. Next the pellet was resuspended in FACS buffer (0.5 % w/v bovine serum albumin (BSA) in PBS, sterile filtered) and endothelial cells were incubated with 1:200 diluted, FITC-conjugated mouse anti-human CD31 (#560984; BD biosciences, USA) and 1:100 diluted, PE-conjugated mouse anti-human CD140b/PDGFRβ (#558821; BD biosciences, USA) for 1 h at 4°C. The pericytes were incubated with 1:100 diluted, PE-conjugated mouse anti-human CD140b/PDGFRβ and 1:500 diluted, anti-NG2 primary rabbit antibody (#ab255811; Abcam, USA). For pericytes, after washing thrice with FACS buffer, 1:500 diluted, AF647-labeled donkey anti-rabbit secondary antibody (A32795, Thermo Fisher Scientific, USA) was added for 1 h at 4°C. Next, cells were washed thrice with FACS buffer and analyzed using FACS Canto II (BD Biosciences, USA).

### Alveolus-on-chip platform design, fabrication and pre-treatment

The alveolus-on-chip platform was adapted from a previously described microfluidic organ-on-chip (OoC) device, consisting of an open-top gel chamber flanked by two parallel medium channels separated by micro-pillars (**Fig. 1**)[16]. Briefly, the device consists of nine open-top OoC units, each comprising a central gel chamber flanked by two parallel medium channels separated by micro-pillars. The micropillars allow loading of a cell-laden hydrogel into the gel chamber without leaking into the medium channels. The platform has a standard well-plate footprint and is fabricated by micro-CNC milling of a polymethylacrylate (PMMA) block, followed by sealing with an adhesive tape bottom layer. Each OoC unit is independently accessible, enabling independent perfusion and culture. Further fabrication details have been reported previously [16].

**Figure 1.**
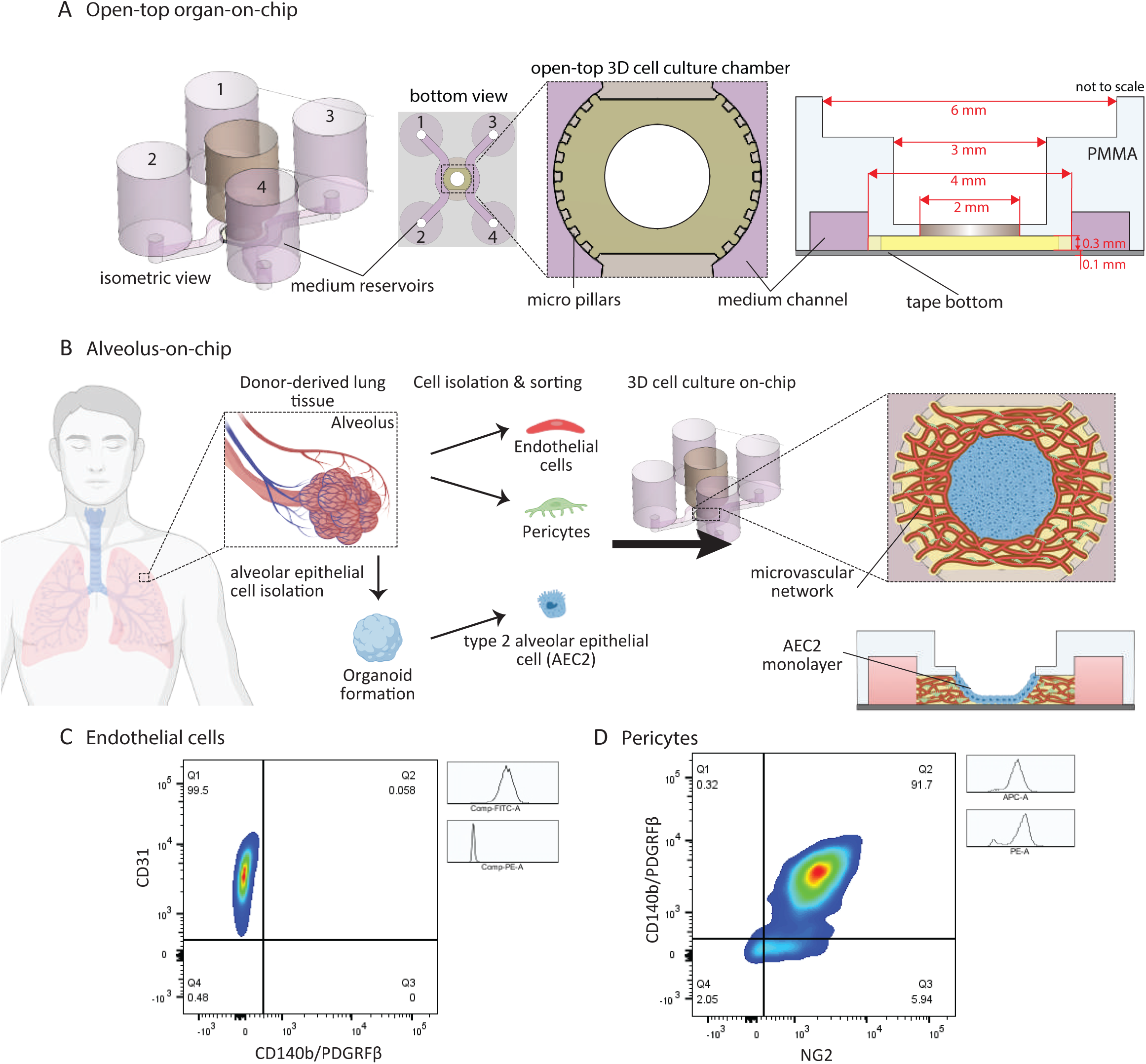
Chip design and characterization of endothelial cells and pericytes isolated from human lung tissue. **(A)** Schematic illustration of the open-top organ-on-chip system, with a bottom side consisting of an open-top gel chamber connected to medium channels through an open row of pillars. Top side consists of medium reservoirs at each end of the medium channels and a separate central medium reservoir surrounded by micro pillars connecting to the open-top gel chamber. **(B)** Experimental set-up alveolus-on-chip: human-derived lung tissues were enzymatically and mechanically dissociated and endothelial cells, pericytes and type 2 alveolar epithelial cells (AEC2) were isolated. To establish vascular networks in the alveolus on-chip, a hydrogel solution is mixed with endothelial cells and pericytes and cultured in the chip to allow formation of vascular networks. A monolayer of AEC2 were cultured on top of the vascular bed. **(C)** Representative flow cytometry analysis of endothelial cells isolated from lung tissue and expanded at passage 1 (P1) for CD140b/PDGRFβ and CD31 (n=4 different donors). **(D)** Flow cytometry analysis of pericytes isolated from lung tissue and expanded at P1 for NG2 and CD140b/PDGRFβ (n=1).

Prior to hydrogel loading the device was surface treated with air plasma (50 W, 45 sec, Cute, Femto Science, South Korea) for further sterilization and to enhance the hydrophilicity of the gel chamber. To enhance adhesion of the hydrogel to the surface of the gel chamber, devices were coated with polydopamine (PDA) as described earlier [17]. Briefly, 100 μl of a sterile filtered 2 mg/ml dopamine hydrochloride solution (Sigma Aldrich, USA) prepared in 10 mM Tris-HCl (pH 8.5) was injected in the gel chamber and incubated for 1 h at RT. The dopamine solution was aspirated, and chips were washed two times with deionized water. The device was fully dried, and the gel chamber immediately loaded with hydrogel.

### Seeding cells in the alveolus-on-chip platform

When confluent, endothelial cells or pericytes were detached using TrypLE^TM^ Select (#12563011; Thermo Fisher Scientific, USA) for 5 min at 37°C. A cell suspension with a volume of 10 µl containing 100,000 endothelial cells and 10,000 pericytes was prepared per chip. To this suspension rat tail collagen 1 (final concentration: 0.2 mg/ml (#A1048301 Thermo Fisher Scientific, USA)) and thrombin (final concentration: 0.1 U/ml (#T7513; Sigma Aldrich, USA)) was added. Fibrinogen (final concentration: 3 mg/ml (#F8630; Sigma Aldrich, US)) was lastly added to the cell mix and a volume of 3.5 µl of the final cell-hydrogel mix was immediately added into the gel chamber through the open top and incubated at 37°C in humidified air for 20 min to allow hydrogel formation. Then, the reservoirs were filled with endothelial medium supplemented with 50 ng/ml of VEGF (#130-109-386; Miltenyi Biotech, the Netherlands) for 48 h and thereafter medium was replaced three times a week without additional VEGF. A hydrostatic pressure gradient was created by adding 200 µl in reservoirs 1 and 2 (left side) and 100 µl in reservoirs 3 and 4 (right side) and the middle reservoir (**Fig. 1**). The chips were placed on top of a rocking platform (OrganoFlow®, Mimetas, the Netherlands), and tilted back and forth at an interval of 10 minutes to 2 hours at 10°, to allow bi-directional perfusion of the gel chamber.

AEC2 were isolated from the same tissue (obtained as per the guidelines of the ethical statement mentioned above) as the endothelial cells and pericytes, using HTII-280^+^ cell sorting and expanded as organoids as previously described [18]. The AEC2 organoids in Cultrex reduced growth factor Basement Membrane Extract, type 2 (BME2, #3533-001-02, R&D systems, Minnesota, US) were incubated with 1 U/ml of dispase for 30 min to dissolve the gel. After centrifugation for 5 min at 265 *g* the pellet was resuspended in 1 ml of TrypLE^TM^ Select and incubated for 5 min at 37°C to dissociate the organoids into single cells. In each chip, 50,000 alveolar cells were seeded in the middle reservoir, on top of the hydrogel containing the vascularized networks with alveolar seeding medium (#100-0847; PneumaCult^TM^, STEMCELL technologies, Canada (**Fig. 1A**). The cells were allowed to settle for 30 min after which additional 150 µl of alveolar seeding medium was added. After 24h, the seeding medium was replaced with 150 µl of the alveolar expansion medium (#100-0847; PneumaCult^TM^, STEMCELL technologies, Canada) in the middle reservoir.

For whole cigarette smoke (WCS) exposure, AEC2 cells were directly seeded 4 h after loading the cell-laden hydrogel. This timing allowed for the formation of an intact AEC2 monolayer within two days, prior to WCS stimulation, at a time when vascular network formation was still actively ongoing. During WCS exposure, the medium channels were sealed and the medium in the middle reservoir containing AEC2 monolayer was removed and air exposed before WCS exposure. The addition of AEC2 cells in alveolar seeding medium and later with alveolar expansion medium, over the cell-laden hydrogel this early did not affect vascular network formation.

### Immunohistochemistry

Upon harvesting, chip cultures were fixed with 4 % paraformaldehyde (PFA) solution. The PFA solution was added in all the chip channels and middle reservoir and incubated for 1 h at RT, washed twice and stored with PBS with primocin® (#ant-pm-1, Invivogen, USA) at 4°C until further used. For staining, permeabilization buffer (0.3 % Triton-X-100 v/v in PBS) was added to the chip channels (200 µl in the left channel and 100 µl in the right channel and middle reservoir) to create hydrostatic pressure and incubated for 20 min at RT and repeated thrice. This pressure gradient allows the entry of the antibodies into the vascular networks. This procedure was repeated with blocking buffer (3 % BSA, 0.3 % Triton-X-100 in PBS and 0.1% Tween20 in PBS), and incubated overnight at 4°C. Next, the chips were incubated with primary antibodies or isotype controls (Table S1) diluted in antibody buffer (0.1 % BSA, 0.3 % Triton-X-100 in PBS and 0.1 % Tween20 in PBS) and incubated for 3 days at 4°C using the same volumes as described for the permeabilization buffer. The chips were washed thrice with permeabilization buffer for 20 min each at RT, followed by addition of secondary antibodies (Table S1) and 4’,6-diamidino-2-phenylindole (DAPI; nuclear stain), diluted in antibody buffer overnight at 4°C. The next day, after washing three times, the chips were stored in PBS until analysis. Imaging was performed using either 10X or 40X oil objective confocal microscope (Leica Biosystems, Germany), equipped with a Dragonfly® 500 or 200 spinning disk (Andor technology, Oxford Instruments, Belfast, UK).

Vessel diameters were measured to compare vascular endothelial networks with and without pericytes using Image J software. For each confocal image, we selected five regions of interest (ROIs), each measuring 250 µm by 250 µm, ensuring an equal number of vessels were analyzed under both conditions.

### Perfusability assay

The alveolus-on-chip models were established with endothelial cells and pericytes as described above. The formed vascular networks were fixed with 4% PFA and stored as described in the immunohistochemistry section. Next, networks were stained and visualized using Texas red^TM^ phalloidin that binds to filamentous actin in the cells (#T7471, Thermo Fisher Scientific, USA) similar as described above for 4h at RT. The perfusion of the vascularized networks was assessed using a 0.02 % (w/v) suspension of Fluoro-Max dyed blue aqueous fluorescent beads (#B0200B; Thermo Fisher Scientific, USA). These beads comprise 1% solids and are made of polystyrene with a density of 1.05 g/cm^3^. To visualize bead flow through the networks, 200 µL of a 50X bead dilution in PBS was added to the left channel, while 50 µL of PBS was added to the right channel. This created a hydrostatic pressure gradient that drives bead movement from left to right.

Perfusion of monocytes through the vascular networks was assessed using the Tohoku Hospital Pediatrics-1 (THP-1) monocytic cell line (ATCC: TIB-202). These cells were cultured in medium consisting of RPMI 1640 (#22409015, Thermo Fisher Scientific, USA) supplemented with 200 mM L-glutamine (#BE17-605E, Lonza, Switzerland) and 10% fetal calf serum (Bodinco B.V., The Netherlands), in T75 flasks at a density of 0.3 to 0.4 X 10⁶ cells/mL in a total volume of 40 mL. To visualize viable THP-1 monocytes, cells were stained with Calcein-AM (ab141420, Abcam, USA). The Calcein-AM stock was diluted to prepare a 0.2 µM working solution in HBSS (Thermo Fisher Scientific, USA). A total of 1 X 10⁵ THP-1 cells were incubated in 5 mL of the working solution for 1 h at 37°C. After incubation, the cells were centrifuged, washed twice with HBSS, and resuspended in 5 mL of THP-1 medium. A 200 µL aliquot of the fluorescent, live monocyte suspension was then added to the left channels of the chip, while 50 µL of PBS was added to the right channels to establish a hydrostatic pressure gradient for perfusion. The flow of beads and monocytes was recorded using a 10X objective in an EVOS FL Auto Imaging system (Thermo Fisher Scientific, USA) for 5 min or monocytes were recorded using a Leica DMi8 microscope equipped with an Andor Dragonfly 200 spinning disk confocal using a 10× objective (NA 0.30×) for 10-20 sec.

### Gene expression analysis

Gene expression analysis was performed on endothelial cells cultured as monolayers in 2D or as vascularized networks with or without pericytes in hydrogel as 3D for four different donors. For 2D culture endothelial cells at passage 1 were seeded in 6W plates, precoated with 50X diluted Matrigel® (#354230, Corning, USA) using endothelial medium. The vascular networks were formed by seeding endothelial cells with and without pericytes in the chip as described earlier. At day 6, endothelial cells in the 2D monolayers and 3D vascular networks were lysed and total RNA was robotically extracted using the Maxwell® 16 simply RNA tissue kit (Promega, the Netherlands) according to manufacturer’s instructions. Gene expression was analyzed by quantitative PCR, where the relative standard curve method was used to derive gene expression and normalized to reference genes ATP synthase F1 subunit Beta (*ATP5B*) and Ribosomal Protein L13a (*RPL13A*). cDNA was mixed with IQ SYBR green mix (Bio-Rad) and primers (Table S2) and the following PCR protocol was performed: started with 95°C for 3 min, followed by 40 cycles for 5 sec at 95°C and 30 min at 63°C, followed by the generation of a melt curve using a gradient from 65°C to 95°C. Data analysis was performed using CFX Maestro software (Bio-Rad, the Netherlands). The endothelial cells in 2D and 3D were analyzed for the following markers, endothelium (*CD31*), macrovascular endothelium (*LYVE1*), microvascular endothelium (*CLDN5*), gCAP microvascular endothelium (*PTPRB, GPIHBP1*), and aCAP microvascular endothelium (*EDNRB1*). The primer pairs used for gene expression analysis are provided in Table S2.

### Whole cigarette smoke exposure on the chip platform

The open-top alveolus-on-chip was exposed to either whole cigarette smoke (WCS) or normal air as a control in a similar fashion as previously reported for cell culture inserts, however now performed at RT [19]. The left and right channels of the chip were sealed with tape (13 mm thickness, Tesa®) to expose only the middle reservoir and prevent WCS/air from entering the other channels. Before WCS exposure, medium in the middle reservoir containing AEC2 monolayer was removed to allow WCS exposure onto the cells. WCS from one cigarette burning over the course of time of 4-5 min was infused in the exposure chamber containing the chip cultures and exited the chamber through a pre-weighed filter at a rate of 1 L/min. Residual smoke was subsequently removed by ventilating room air into the chambers for another 10 min. Air-exposed cell cultures underwent the exact same procedure, however, in separate chambers that were only infused with normal air. After exposure, chip cultures were washed with PBS and endothelial medium was replaced in the left and right reservoirs. The filter was weighed and the total particle matter per liter was calculated as a control measure between smoke exposures.

## Results

### Chip set-up and characterization of cells isolated from lung tissue

The microfluidic chip design that was used to establish a model of the alveolar-endothelial interface, is based on an open-top gel chamber in which the co-culture of primary vascular networks in close contact with a primary alveolar epithelial monolayer is established. This chamber is flanked by two rows of micropillars on both sides to retain the hydrogel in the middle chamber. The chamber is connected via the rows of micropillars to two medium channels and enables medium perfusion through the chamber. The medium channels each have reservoirs on top of their inlets and outlets, respectively. The full schematic of the chip design is depicted in the horizontal and vertical view (**Fig. 1A**). To support our strategy of culturing a vascularized hydrogel containing endothelial cells and pericytes with a monolayer of type 2 alveolar epithelial cells (AEC2) on top of the vascular bed, we isolated pulmonary endothelial cells, pericytes and AEC2 from human lung tissue (**Fig. 1B**). After processing the tissue and generating a single cell suspension, endothelial cells, pericytes and AEC2 were sorted and characterized for their respective cell surface markers using flow cytometry analysis before seeding cells in the chip. At passage 2, nearly all expanded endothelial cells were positive for the expression of CD31 and negative for PDGFRβ, a marker for endothelial-to-mesenchymal transition (**Fig. 1C**). When measured for several donors (n=4), the expanded endothelial cells were ∼97% ± 3.2% (mean ± standard deviation) positive for CD31 expression (Fig. S1A). The morphology of the monolayers of the lung endothelial cells was uniform as shown by the bright field microscopy images (Fig.S1B). The pericytes expanded at passage 2, were ∼92% double positive for both NG2 and the mesenchymal marker, PDGFRβ ([20]; **Fig. 1D**). To reduce the impact of donor-to-donor variability, epithelial cells and endothelial cells were used from various donors, while the pericytes used in all the experiments were derived from the lung tissue of one non-COPD donor.

### Vascular network formation

Next, we assessed if primary pulmonary endothelial cells could form vascular networks in our chip platform. We hypothesized that pericytes may be required for vascular network formation. To test this hypothesis, endothelial cells alone or combined with pericytes were embedded in a fibrin-collagen hydrogel in the chip platform. Bi-directional flow in the chip was achieved by placement on a rocking platform, to establish a hydrostatic pressure gradient to drive the flow of medium through the hydrogel and promoting the self-organization and formation of vascular networks (**Fig. 2A**). We first compared network formation in the presence and absence of pericytes at day 12. In the presence of pericytes, thinner vessel size was observed (**Fig. 2B**), whereas in the absence of pericytes a thicker vessel size was observed (**Fig. 2C**). The vessel diameter quantified in different regions of the chip confirmed that endothelial networks with pericytes had significantly lower vessel diameters compared to networks established without pericytes (**Fig. 2D**).

**Figure 2.**
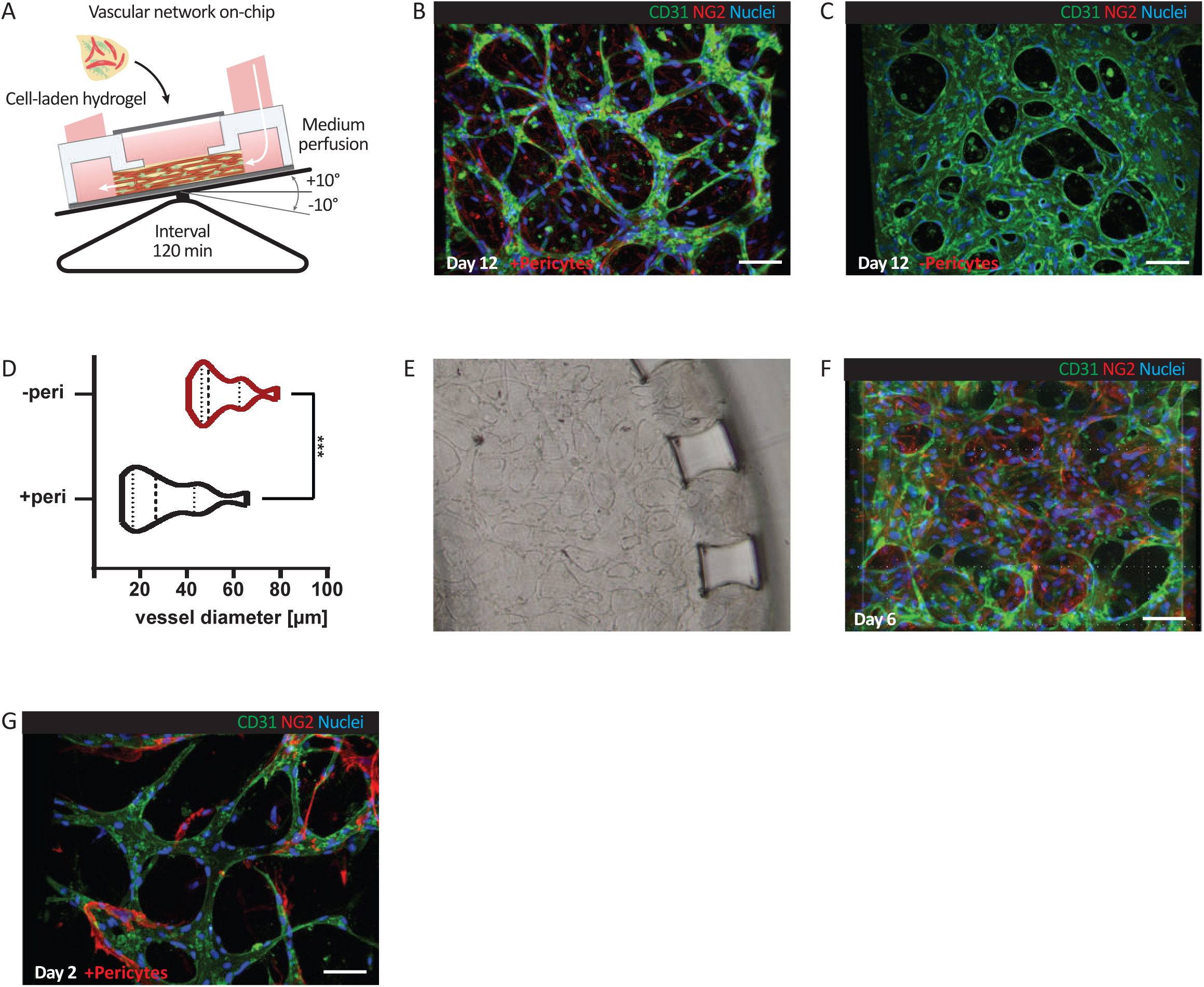
Vascular network formation using human-derived primary endothelial cells and pericytes on-chip. **(A)** Schematic representation of the experimental set-up, with the chip loaded with a cell-laden hydrogel containing endothelial cells and pericytes and medium added to all reservoirs. The chip platform is placed on a rocking platform on a 10° angle and hydrostatic pressure is generated by moving backward and forward at an interval of 2 hours. This allows perfusion of the gel-chamber that promotes vascular network formation. Immunofluorescence images of vascular networks at Day 12 with **(B)** and without **(C)** pericytes with endothelial cells in green (CD31), pericytes in red (NG2) and nuclei in blue (DAPI) Scale bars: 50 µm. **(D)** Violin plot comparing vessel diameters between conditions with (+peri) and without (-peri) pericytes, showing a significant decrease in vessel diameter with pericyte presence (p < 0.001). Analysis of differences was conducted using unpaired two-tailed t-test. **(E)** Representative bright field image (n=3 different donors) after 6 days of self-assembled vascular network on-chip. Scale bar, 100 µm. **(F)** Representative confocal image (n=3 different donors) after 6 days of a self-assembled vascular network formation on chip with endothelial cells in green (CD31), pericytes in red (NG2), and nuclei in blue (DAPI). Scale Bar, 50 µm. **(G)** Immunofluorescence image of early vascular network formation at day 2 with pericyte interaction with vessels. Scale bar: 50 µm.

Since the alveolus comprises a microvascular network with vessel size of ∼10 µm in diameter [21], the smaller vessel diameter in the presence of pericytes is approaching this aim and we therefore continued with the endothelial/pericyte co-culture in the chip. Already on day 6, a continuous, self-assembled, endothelial-pericyte vascular network was formed, as demonstrated with bright-field microscopy (**Fig. 2E**). Next, we confirmed preservation of endothelial phenotype in the 3D vascular networks with analysis of CD31 expression (**Fig. 2F**). Of note, we also confirmed that pulmonary endothelial cells isolated from two different COPD patients (stage II and stage IV) were able to form vascular networks (Fig. S1C). However, we did observe that vascularization of the endothelial cells is donor-dependent and using cells of ∼18% of the donors (independent of COPD status) no network formation could be observed, in our hands. Also, when mesenchymal cells were not removed efficiently during endothelial cell isolation, the morphology of cells in the monolayers was altered (Fig. S1D), with a decrease in CD31 expression and increase in mesenchymal cell marker, PDGFRβ (Fig. S1E). These cells were also not able to form vascular networks when co-cultured with pericytes (data not shown).

The location of the pericytes within the networks was visualized by pericyte-specific NG2 expression. Image analysis confirmed the presence of 3D-vascular networks with CD31-expressing endothelial cells and NG2-expressing pericytes (**Fig. 2F** and Video S1). We also tested for periaxin, a protein associated with lung microvascular endothelial cells [22]. However, the pulmonary endothelial cells in both vascular networks or monolayers did not express periaxin (data not shown).

Pericyte distribution throughout the hydrogel was not random, but occurred in close proximity to the formed endothelial vascular networks (Fig. S1F). To further study endothelial-pericyte interactions, we characterized the network formation at an earlier time point, as network formation starts at 48 hours after cell seeding. On day 2, the pericytes, marked by NG2, were already in close proximity to the endothelial cells, wrapping around the CD31^+^ endothelial vessel-like structures (**Fig. 2G**). Overall, this demonstrates that for in vitro vascular network formation, integration of pericytes is relevant, they interact with the endothelial cells, can maintain their phenotype, and alter the vascular endothelial network formation in our chip platform.

### Characterization of cells cultured in the vascular network

To characterize the phenotype of endothelial cells cultured in vascular networks, we performed two sets of analyses. First, we analyzed gene expression patterns of endothelial cells cultured in 2D monolayers and compared them to gene expression in 3D vascular networks at day 6. As pericytes could not be cultured within the 2D endothelial monolayers, we established 3D endothelial networks without pericytes for a fair comparison. Results showed that endothelial cells cultured in 3D networks were significantly enriched for *CD31* expression compared to endothelial cells cultured in 2D monolayers. Expression of genes associated with various endothelial subsets [22, 23], i.e. macrovascular (*LYVE1*), microvascular (*CLDN5*), general capillary (gCap; *GPIHBP1*, *PTPRB*), aerocyte capillary (aCap; *EDNRB1*) and alveolar repair (*BMP6*) was not significantly different between the two groups (**Fig. 3A**).

**Figure 3.**
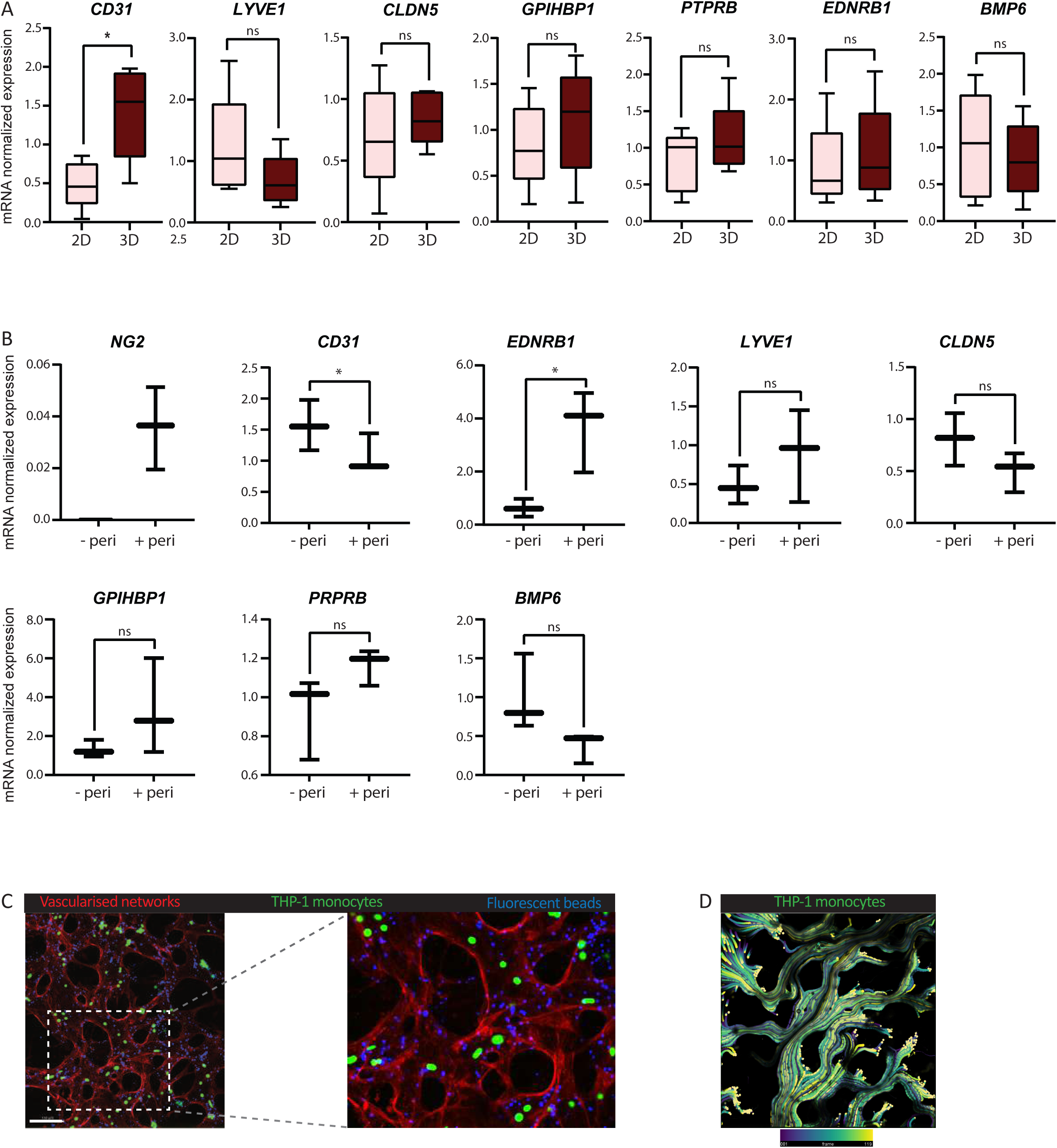
Characterization of vascular network formation on-chip. **(A)** qPCR analysis of mRNA expression of genes expressed in endothelial cells (*CD31*), macrovascular endothelium (*LYVE1*), microvascular endothelium (*CLDN5*), gCAP-microvascular endothelium (*GPIHBP1*, *PTPRB*), aCAP-microvascular endothelium (*EDNRB1*) and alveolar repair (*BMP6*) in endothelial monolayers (2D) versus endothelial networks without pericytes (3D) (n=4 different donors). Box plots display normalized mRNA expression levels. Analysis of differences was conducted using paired two-tailed t-test. Statistical significance is indicated by p < 0.05 (*), while "ns" denotes non-significant differences. Data represent mean ± interquartile range. **(B)** qPCR analysis of mRNA expression of markers for endothelial cells (*CD31*), macrovascular endothelium (*LYVE1*), microvascular endothelium (*CLDN5*), gCAP-microvascular endothelium (*GPIHBP1*, *PTPRB*), alveolar repair marker (*BMP6*), pericyte marker (*NG2*) and aCAP-microvascular endothelium (*EDNRB1*) in endothelial vascular networks with and with pericytes (n=3 different donors). Analysis of differences was conducted using paired two-tailed t-test. Statistical significance is indicated by p < 0.05 (*), while "ns" denotes non-significant differences. Normalized gene expression was calculated using the reference genes ATP synthase F1 subunit Beta (*ATP5B*) and Ribosomal Protein L13a (*RPL13A*). **(C)** Representative fluorescent image of self-assembled vascular network (red - phalloidin) perfused using fluorescent beads (blue, 2 µm diameter) and live THP-1 monocytes (green). Scale bar, 150 µm. **(D)** Flow of the calcein-AM-labeled THP-1 monocytes (viridis) through the vascular networks were recorded for 20s and converted to flow trajectories using ImageJ. Colors depict frames; dark blue (frame 1) to yellow frame 119. Scale bar, 200 µm.

Next, we tested if incorporation of pericytes into the vascular networks affected endothelial cell phenotype, and compared endothelial cells cultured in 3D networks in presence or absence of pericytes at day 6. As expected, the incorporation of pericytes into the vascular endothelial networks resulted in the expression of the pericyte marker *NG2*. We observed a relative reduction in *CD31* expression in the co-culture, which is likely due to the fact that pericytes do not express *CD31* but do contribute to normalization of the total expression. Interestingly, gene expression of the aerocyte capillary marker *EDNRB1* was significantly higher in networks cultured with pericytes compared to those without pericytes (**Fig. 3B**), even though pericytes do not express this gene (**Fig. 3A**). This suggests that the observed increase is due to enhanced expression by endothelial cells. Expression of genes associated with other endothelial subsets, i.e. *LYVE1, CLDN5, GPIHBP1, PTPRB* and *BMP6* were not significantly different between the two groups (**Fig. 3B**). A 2D monolayer of pericytes did not express any of the above mentioned endothelial markers including *CD31*, *LYVE1, CLDN5, GPIHBP1*, *PTPRB, BMP6* and *EDNRB1* that were expressed in endothelial cells and therefore it is highly unlikely that the pericytes would contribute to the detected expression in the co-cultures. However, the pericytes did express their surface marker *NG2*, which was absent in the endothelial monolayer (Fig. S2A) Finally, we tested if MRC5 fibroblasts were also able to support endothelial vessel formation, like pericytes. The networks observed on day 6 showed that the addition of MRC5 fibroblasts (marked by α-SMA) resulted in disrupted endothelial networks (Fig. S2B). However, the presence of pericytes (in this case also marked by α-SMA) allowed continuous endothelial network formation (Fig. S2C). Fibroblasts were therefore not as potent in supporting vascular network formation as pericytes.

### Perfusion of the engineered vascular networks

To determine whether the established vascular networks could be actively perfused from one side of the chip to the other side, we tested their perfusion with fluorescent beads and immune cells. When the endothelial-pericyte networks were well-established by day 6, the vascular networks were either directly perfused with fluorescent beads (Video S3), or fixed and labeled with phalloidin to visualize the networks before perfusion with monocytes. The phalloidin-stained vascular endothelial networks enabled tracking of the fluorescent beads and monocytes within the borders of the endothelial networks. Fluorescent beads (2 μm in diameter) were introduced in the medium channel reservoirs and bead locations were tracked over time. In addition, calcein-AM fluorescent THP-1 monocytes were also introduced to assess if immune cells, which are larger in size than the used beads, could also move through the networks. The beads and monocytes were distributed throughout the networks within seconds after introduction to the chip inlet, and exited the networks on the other side, demonstrating that the formed endothelial networks were capable of immune cells perfusion (**Fig. 3C** and Video S4). Monocytes were imaged for max. 20 seconds and cell trajectories were visualized based on a time-resolved intensity projection of the fluorescent signal (**Fig. 3D**). If network formation was suboptimal, the beads stayed immobile in between the pillars on the entry side and were unable to move through the gel chamber (Fig. S3). Altogether, the endothelial-pericyte vascular networks are capable of perfusing small particles and live immune cells.

### Culture of an AEC2 monolayer on top of endothelial-pericyte vascular networks

In the alveoli, the epithelial cells are in close proximity to the endothelial cells. Previously, we showed that a monolayer of pulmonary endothelial cells supports the formation of type 2 alveolar organoids in a static insert model [3]. For the alveolus-on-chip, we therefore next cultured primary AEC2 on top of the endothelial cells and pericytes cultured in a hydrogel. On day 5 after the networks were formed, an AEC2 suspension was seeded on top and allowed to form a monolayer, as depicted in the schematic representation (**Fig. 4A**). The compatibility of co-culturing these cells in the chip system was tested until day 12. Image analysis of the chip demonstrated the presence of an intact AEC2 monolayer that is marked by pro-surfactant protein C (SFTPC) expression which is surrounded by endothelial networks marked by CD31 expression (**Fig. 4B-C** and Video S2). Interestingly, we observed that AEC2 seemed to have a variable expression level of pro-SFTPC, with higher expression in cells close to the endothelial networks compared to cells in the middle. Here, we demonstrated that endothelial vascular networks were successfully co-cultured with an AEC2 monolayer using lung tissue-derived cells in the alveolus-on-chip platform.

**Figure 4.**
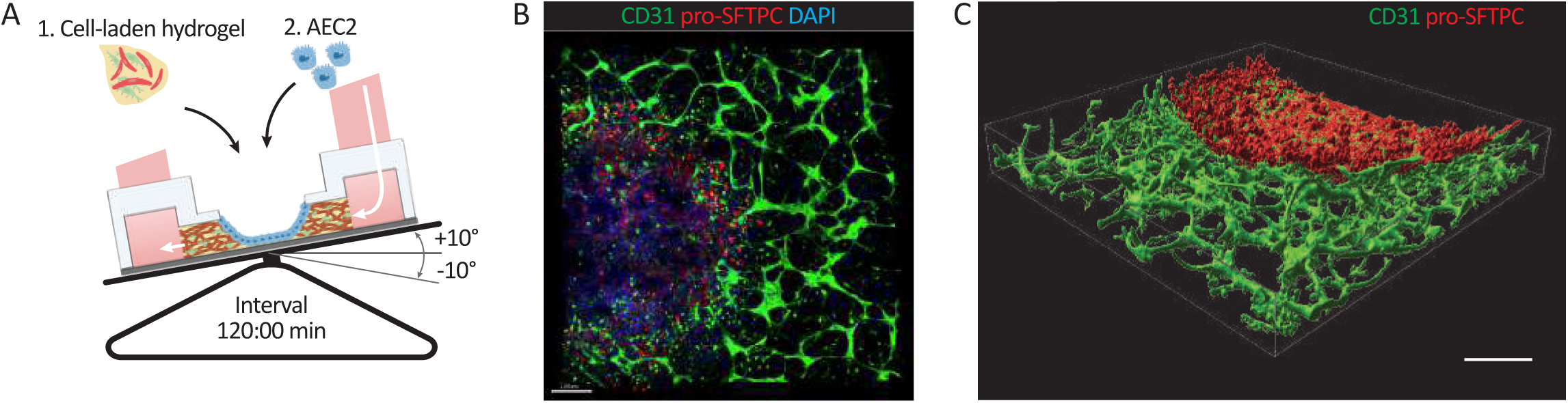
Alveolus on-chip model with AEC2 monolayer co-cultured on self-assembled microvascular network. **(A)** Schematic representation of experimental set-up with cell-laden hydrogel containing endothelial cells and pericytes in the open-top area. After seeding, the chips were mounted on a rocking platform at a 2 hour interval on a 10° angle. After formation of vascular network at day 5, AEC2 were seeded into the microwell to form a monolayer with direct interaction. **(B)** Representative confocal image (n=3 different donors) of alveolus on-chip with self-assembled vascular network stained for CD31 (in green) and AEC2 monolayer stained for pro-SFTPC (in red) and nuclei stained with DAPI (in blue). Scale bar: 20 µm. (Left panel). **(C)** 3D reconstruction of the alveolus-on-chip, with the AEC2 monolayer (red) and the microvascular network (green). Scale bar: 500 µm (right panel).

### Endothelial-pericyte networks are more susceptible to damage by cigarette smoke in presence of alveolar epithelial cells

With its open-top form, the chip platform is highly suitable for studies with air-borne substances. We therefore tested if the alveolus-on-chip cultures could be successfully exposed to whole cigarette smoke (WCS). Endothelial cells and pericytes were seeded in the fibrin-collagen hydrogel and after 4 hours AEC2 were seeded on the hydrogel. In these experiments, the AEC2 were seeded on the same day as the endothelial cells to allow formation of an intact monolayer within two days, ensuring that only the AEC2 were directly exposed to WCS exposure and not the vascularized networks. Two days after establishing the co-culture, AEC2 monolayers and vascular networks were established, and chips were exposed to WCS once daily for five consecutive days and were analyzed 24 h later (**Fig. 5A**). The chips were exposed to WCS or air as control, in their respective chambers, for 15 min with the alveolar medium completely removed from the inner well. The WCS was thereby in direct contact with the AEC2 monolayers and not with the vascular networks (**Fig. 5B**). After harvesting the chip cultures, they were stained for CD31 and NG2 (markers for endothelial cells and pericytes, respectively). The AEC2 monolayer was stained with pro-SFTPC. Analysis showed that, in air-exposed controls, continuous networks were present, indicated by expression of CD31^+^ endothelial cells and NG2^+^ pericytes (**Fig. 5C**). Repeated WCS exposure completely disintegrated endothelial-pericyte networks, indicated by non-continuous vessels and presence of small clumps of CD31^+^ cells. However, the AEC2 monolayer, directly exposed to WCS, remained intact with no change in morphology compared to AEC2 monolayers that were exposed to air (**Fig. 5C** and **Fig. 5D**; the damage to the epithelial layer in the top right of the respective images was due to handling during staining procedure). Next, we analyzed if the disruption of vascular networks by WCS exposure was mediated by AEC2. To this end, vascular networks were only exposed to WCS, in a similar fashion as the previous set-up, in presence or absence of an AEC2 monolayer. In the absence of AEC2, upon repetitive WCS exposure, the endothelial-pericyte vascular networks remained intact with no signs of disruption (**Fig. 5E**). However, WCS-exposed chip cultures with an intact AEC2 monolayer showed again a prominent loss of vascular networks (**Fig. 5F**), suggesting that the AEC2 cells are involved in the vascular damage upon exposure to WCS. Together our results showed the feasibility of whole cigarette exposure to the alveolus-on-chip platform and the involvement of AEC2 in vascular damage in response to WCS exposure.

**Figure 5.**
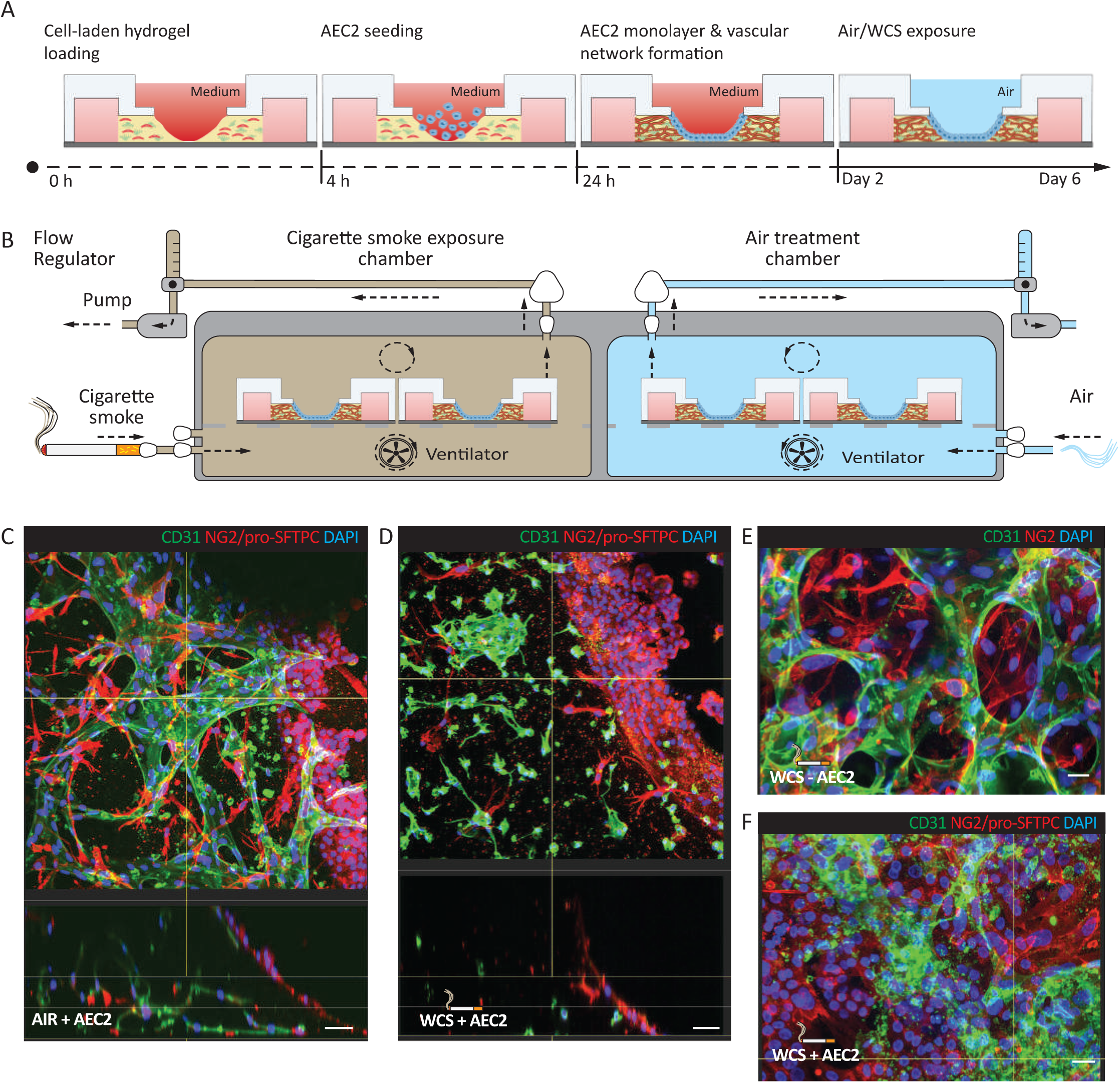
Effects of whole cigarette smoke exposure on vascularized endothelial networks in co-culture with AEC2 monolayer. **(A)** Schematic representation of the experimental set-up of whole cigarette smoke exposure on alveolus on-chip. Endothelial cells and pericytes were embedded in hydrogel and after 4 hours AEC2 cells were seeded as a monolayer over the hydrogel. The vascular networks self-assembled within 48 hours and an intact AEC2 monolayer was formed. Starting at day 2, the AEC2 monolayer was exposed to either whole cigarette smoke (WCS) or air every day, until day 5 and analyzed at day 6. **(B)** Schematic representation of whole cigarette smoke and air chamber. The chip cultures were exposed to WCS by placing them in an exposure chamber infused with WCS or air for 4 to 5 min and the chambers were ventilated with room air for 10 min. Representative confocal image (n=3 different donors) of air-exposed alveolus-on-chip with continuous vascular network **(C)** and WCS-exposed alveolus-on-chip with disrupted vascular networks **(D)** marked with endothelial marker, CD31 (in green), pericyte marker, NG2 (in red) and AEC2 monolayer in the middle stained for pro-SFTPC (in red). Scale bar, 25 µm. **(E)** Representative confocal image (n=2 different donors) of WCS-exposed chip with vascular networks and without an AEC2 monolayer marked by CD31 (in green) and NG2 (in red). **(F)** Representative confocal image of co-cultured vascular networks marked by CD31 (in green) and AEC2 monolayer marked by pro-SFTPC (in red). Scale bar, 10 µm.

## Discussion

In our open-top, alveolus-on-chip model, we have generated self-assembled, 3D-vascularized stable and perfusable networks consisting of primary human endothelial cells and pericytes, co-cultured with alveolar epithelial cells for up to 7 days. This chip model employs hydrostatic pressure to create interstitial flow through the hydrogel, promoting vascular network formation. Vessel formation starts from 48 hours after seeding endothelial cells and pericytes, with perfusable networks observed within 6 days, although this timing was highly donor-dependent. The vascular network structure can be maintained for at least 12 days. Furthermore, we demonstrate that the integration of pericytes alters network morphology, including vessel diameters. Close contact between endothelial cells and pericytes was already observed from day 2 onward, supporting the view that pericytes, normally present in human alveolar vascular beds [24], play a key role in the formation of capillary-like sized vessels. Integration of pericytes is crucial in our chip cultures, as they support the development of microvascular-like size vessels, contribute to the maintenance and stability of the vasculature [25] and are involved in alveologenesis [26]. The ratio of endothelial cells to pericytes in the lung is estimated to be around 9:1 [27], and therefore we used similar ratios to be as comparable as possible. Studies have shown that the morphology of pericytes can vary along the capillary wall [28]. However, we were not able to observe differences in morphology of pericytes attached to vessels of different diameters.

The development of a vascular network in a fibrin hydrogel is not novel, since vessel-on-a-chip models have shown that this is possible with a wide variety of cells, also including human induced pluripotent stem cell-derived endothelial cells [29, 30] and human umbilical vein endothelial cells (HUVECs) [31]. However, we have used primary human lung microvascular endothelial cells, and additionally we demonstrated the feasibility of culturing a monolayer of AEC2 directly on the vascularized hydrogel, to more closely mimic the local environment in the alveolus. Importantly, our platform facilitates the study of direct interactions between human primary alveolar epithelial cells and vascular endothelial networks by eliminating the porous membrane typically used to separate cell compartments in conventional lung-on-chip models [32, 33]. This membrane-free configuration enables more physiologically relevant cell culture, while still supporting the establishment of an air-liquid interface (ALI), a critical feature for mimicking the lung environment. In the present study, ALI was maintained only transiently during air or WCS exposures. Further model development is required to sustain ALI over extended periods. One other report includes a co-culture of a vascular network with alveolar epithelial cells, however for this study it seemed like a single donor (different donors for different cell types) was used, cells were commercially sourced and the perfusability of the vasculature was not described [34]. Here, we have demonstrated the feasibility of using several donors and donor combinations and patient-derived endothelial cells to generate lung vascular networks. In future studies, sourcing cells (endothelial cells, pericytes and AEC2) from lung tissue of the same donor would allow more personalized disease modelling for e.g. drug testing and would allow comparison between diseased cells and control cells. In our preliminary data, endothelial cells isolated from COPD donors and non-COPD donors had comparable network formation time when cultured with non-COPD pericytes. We hypothesize that the COPD-specific features in endothelial cells [35] may be normalized during expansion of endothelial cells for example because they arise from only a small proportion of the isolated CD31^+^ cells that are viable. Alternatively, it could be that a COPD-specific phenotype in the endothelial cells is masked by the presence of healthy pericytes. Follow-up studies will be needed to further characterize the phenotype of the endothelial cells from healthy and diseased donors, as well as systematically combining endothelial cells and pericytes from different donors.

We also demonstrate that endothelial cells within vascularized networks have increased *CD31* gene expression compared to endothelial monolayers, suggesting a better preservation of endothelial phenotype and potentially better cell-cell adhesion. In the lung, several endothelial cell subsets including lymphatic, arterial, venous, capillary endothelial cells, have been identified [22]. The networks in our chip system were negatively stained for periaxin, a protein described to be enriched in capillary endothelial cells in the alveolus [22], despite the fact that expression of the general microvascular marker *CLDN5* was readily detected. Endothelial cells isolated and expanded from lung tissue as 2D monolayers were also negative for periaxin (data not shown), which indicates that periaxin expression could be diminished after isolation or expansion already. Importantly, the addition of pericytes to the endothelial vascular networks was found to increase gene expression of *EDNRB1,* which is shown to be enriched in capillary microvascular endothelial cells, present in the alveolar compartment [22]. Two capillary endothelial subsets have been identified based on scRNAseq analyses, the general capillary endothelial cells (gCap) and aerocyte (aCap) endothelial cells, which closely interact with AEC2 and alveolar type 1 cells (AEC1), respectively [23]. The cells in the endothelial networks in our chip did express gCap and aCap alveolar capillary markers. Further research is required to determine whether culturing AEC2 or AEC1 monolayers on the network can promote the formation of gCAP or aCAP endothelial cell subsets. During alveolar injury, AEC2 act as progenitor cells that can differentiate into AEC1 [36], which are required for gas exchange. In the current setup, we showed the compatibility of culturing AEC2 monolayers on lung vascular networks comprised of endothelial cells and pericytes. However, work on a controlled differentiation on the chip of AEC2 to AEC1 is still underway and would further improve human tissue-relevance of the chip culture.

A strength of the current platform is the open-top configuration that allows whole cigarette smoke exposure of the AEC2 monolayer. This enabled us to establish that repeated WCS exposure of the AEC2 monolayer leads to complete disintegration of the vascularized endothelial networks, a process likely mediated by AEC2 cells as vascular networks remained intact in response to WCS in absence of AEC2. In our previous study [3] and in work of others [37], cigarette smoke extract was used to study its effects on lung microvascular endothelial cells. This extract provides a pro-inflammatory stimulus and includes the soluble compounds of WCS, but does not fully recapitulate the more complex exposure of whole cigarette smoke. The open-top design of our chip model enables us to study the effect of airborne WCS, more closely modeling the exposure of the alveoli in smokers. Here, we demonstrated that endothelial cells were more susceptible to cigarette smoke exposure compared to AEC2 as vascular networks disintegrated while the alveolar epithelium remained intact. This is in line with our recent study [3], where we showed that microvascular endothelial cells, but not cells with AEC1 phenotype, were specifically reduced in lung tissue of emphysematous COPD patients compared to non-COPD controls, in sections of comparable volume (relatively unaffected regions). Also *in vivo,* mouse models have shown that cigarette smoke exposure resulted in fewer endothelial cells and pericytes [38]. In our model, exposure of vascular networks to WCS did not affect the network integrity unless AEC2 were present. Our future work will focus on identifying AEC2-derived factors involved in the disintegration of the vascular networks upon smoke exposure.

Clearly these new developments do not come without limitations. The current platform is not able yet to simulate the cyclic mechanical stretch associated with breathing as this is known to influence cellular behavior, promote differentiation, and enhance the physiological relevance of lung-on-chip systems [12]. This constraint arises from both design limitations and the mechanical properties of the materials used in the fabrication of this platform. As the platform is based on hydrostatic pressure driven flow, established by rocking back and forward, the induced flow was bi-directional as opposed to physiological uni-directional flow in the lung capillaries. Nonetheless, another study has shown that vessel formation on-chip was comparable between uni- and bi-directional flow [25]. Controlling the flow rate in our model is limited to the height of the medium column in the medium reservoirs and the tilting angle. This limits control over shear forces on the cells and ultimately, affects endothelial barrier integrity, which can be influenced by shear stress [39]. Further adaptation of the platform is planned to include airflow, uni-directional medium flow and inclusion of sensors. The incorporation of functional assays to monitor endothelial barrier integrity and permeability would significantly strengthen the platform’s utility in modeling disease states. These readouts are particularly valuable for studying barrier dysfunction, a hallmark of various pulmonary pathologies such as acute respiratory distress syndrome (ARDS), pulmonary edema, and inflammation-driven vascular leakage. Additionally, future developments could include the integration of biosensing technology or real-time monitoring systems to track dynamic tissue responses under physiological and pathological conditions. Such advancements would not only expand the platform’s applicability in fundamental research but also enhance its translational relevance for drug screening and disease modeling.

Finally, with the perfusable networks we were able to demonstrate that monocytes could be perfused and tracked through the vascular networks. This work opens up new avenues for investigating immune cell recruitment to the alveolar lumen upon injury. Our model can be furthermore employed in future studies to investigate the impact of inflammatory cytokines, infections and alarmins, elevated in patients with chronic lung disease [40], on pulmonary vascular network formation and permeability. Various forms of damage-inducing factors may promote impairment in endothelial-pericyte cross-talk leading to vascular permeability during lung injury [41]. Studies showed that immune cells such as neutrophils [42] and CD8^+^ T cells [43] in the alveolar regions are involved in pathogenesis of lung disorders. Using our chip model, we have shown that monocytes can be perfused through the vascular networks, thereby broadening the potential to study immune cell interactions in these models. Furthermore, future studies could focus on incorporating immune cells also on the alveolar epithelial monolayer to understand the role of inflammatory innate immune cells in lung disorders.

Overall, we developed an open-top lung-on-chip model for air-borne exposures with a perfusable and stable endothelial-pericyte vascular network using patient-derived cells. This model deploys a membrane-free AEC2 co-culture, cultured over a vascular bed enabling better modelling of epithelial-capillary interactions during cigarette smoke exposure. Thereby, this lung-on-chip platform can serve as a tool to study lung disorders and eventually screen drug targets.

## Acknowledgements

This work was supported by Lung Foundation Netherlands (grant#: 4.1.19.021), the European Union’s Horizon 2020 research and innovation programme under the Marie Skłodowska-Curie Grant No. 812954, and by the Netherlands Organ-on-Chip initiative, an NWO Gravitation project (024.003.001) funded by Ministry of Education, Culture and Science of the government of the Netherlands.

