## Supplementary Figures for "Modelling cigarette smoke-induced lung vascular dysfunction using an alveolus-on-chip"

### Figure S1

A

B

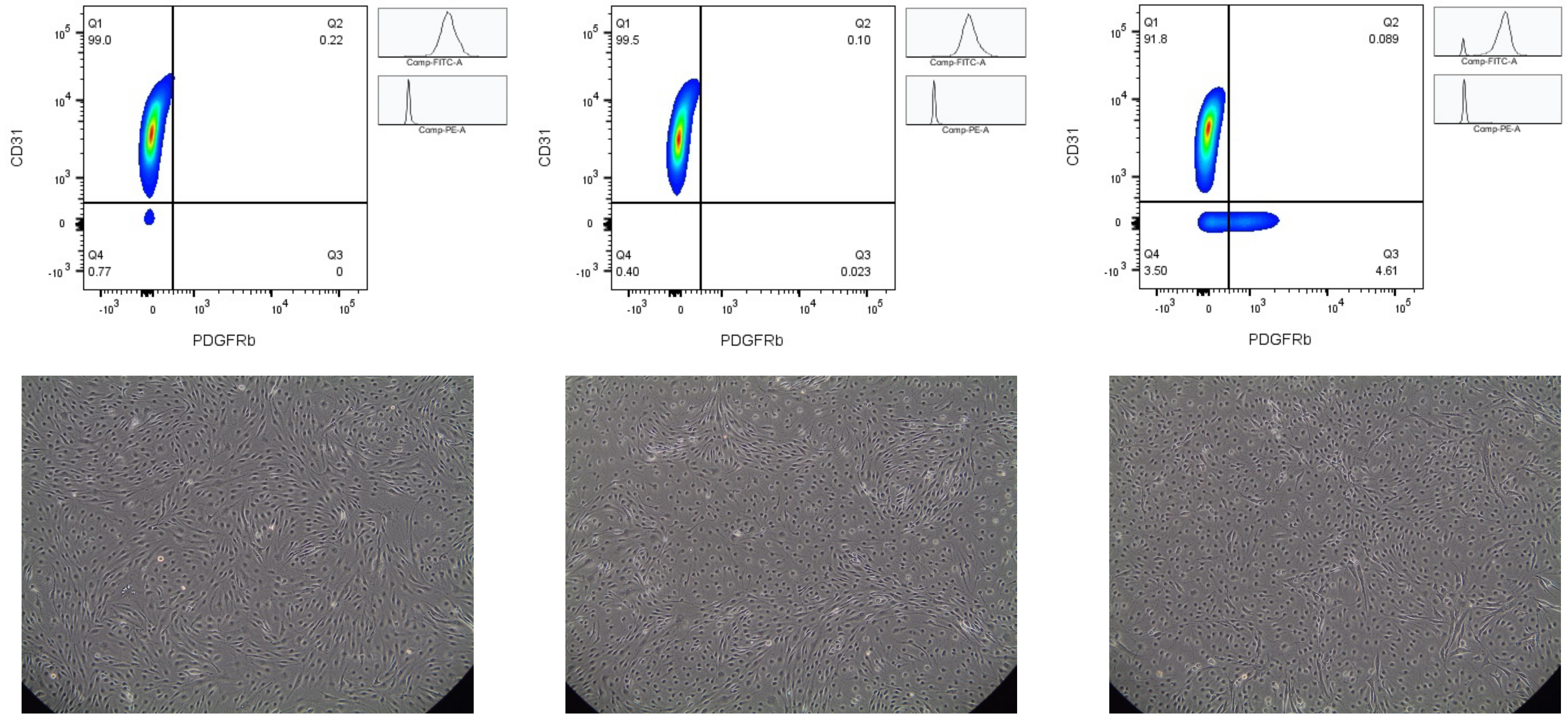


**Figure S1.** (**A**) Flow cytometry analysis of endothelial cells isolated from lung tissue and expanded at P1, for CD140b/PDGRFβ and CD31 (n=3 different donors). (**B**) Bright field images (of n=3 different donors) depicting morphology of endothelial monolayers expanded at P1.

#### Figure S1

C

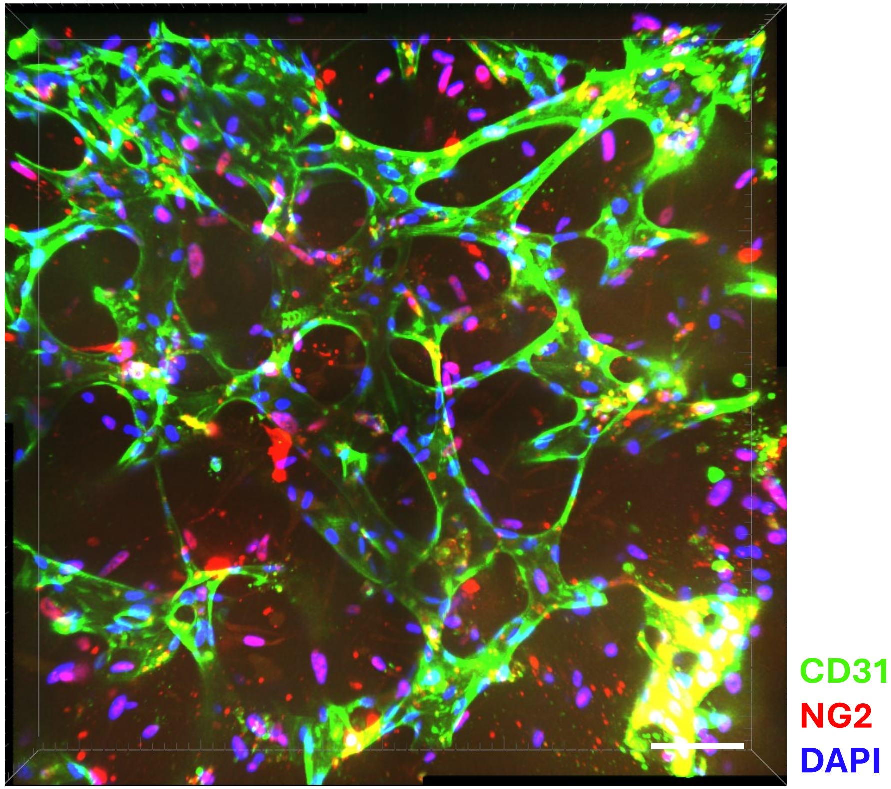


**Figure S1. (C) Vascular network formation of endothelial cells isolated from COPD donor.** Representative confocal image (n=2) after 6 days of a self-assembled vascular network stained with endothelial cell marker, CD31 (in green) and pericyte marker NG2 (in red) and nuclei with DAPI stain (in blue). Scale Bar, 70 µm.

### Figure S1

**D**

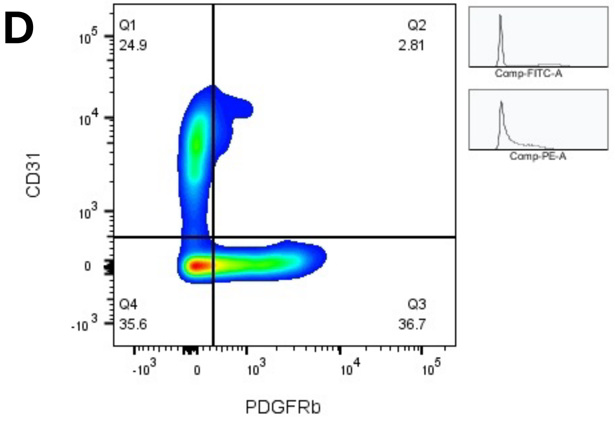


**E**

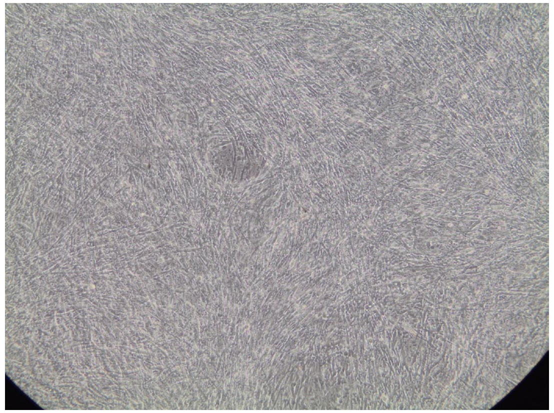

**Figure S1.** (**D**) Flow cytometry analysis of endothelial cells inefficiently isolated with mesenchymal contamination, from lung tissue and expanded at P1, for CD140b/PDGRFβ and CD31 (n=1). (**E**) Bright field images (of n=1) depicting morphology of endothelial monolayers in the presence of mesenchymal cells, expanded at P1.

#### Figure S1

F

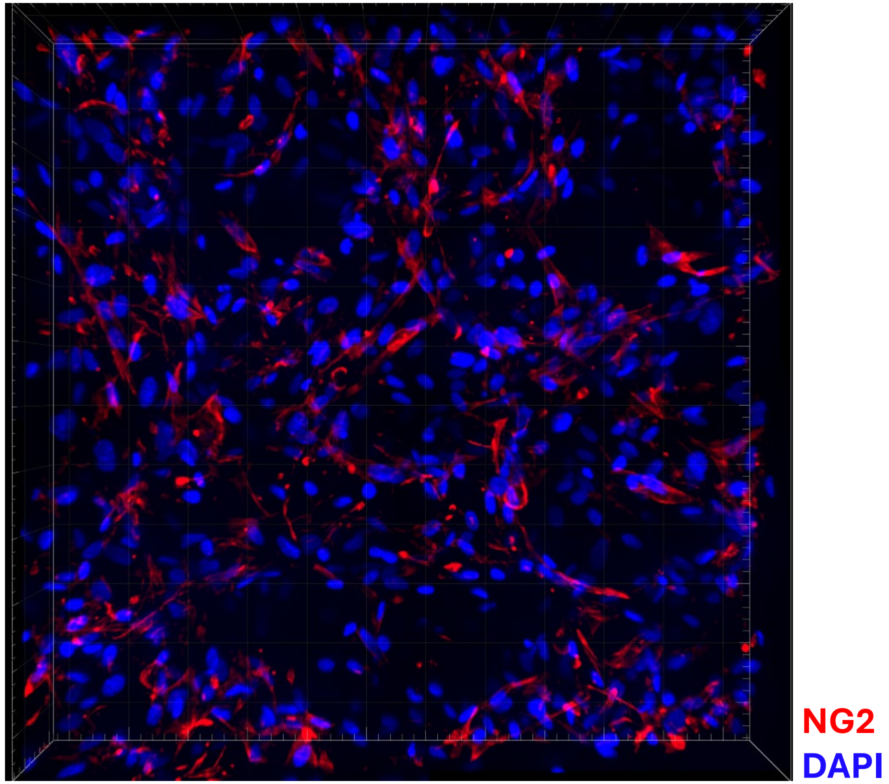


**Figure S1. (F) Pericytes are embedded within the endothelial networks.** Representative confocal image after 6 days of a self-assembled vascular network (n=3 different donors) stained with pericyte marker NG2 (in red) and nuclei with DAPI stain (in blue). Scale Bar, 70 µm.

#### Figure S2

A

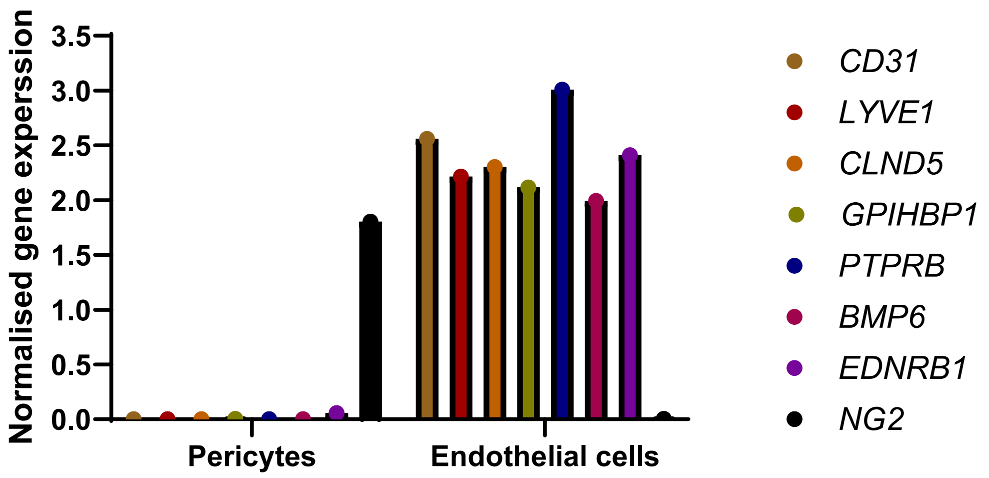


**Figure S2. Gene expression analysis of pericyte monolayer. (A)** qPCR analysis of mRNA expression in pericytes (*n*=1) and enothelial cells (*n*=1) for markers of endothelial cells (*CD31*), macrovascular endothelium (*LYVE1*), microvascular endothelium (*CLDN5*), gCAP-microvascular endothelium (*GPIHBP1*, *PTPRB*), alveolar repair marker (*BMP6*), aCAP-microvascular endothelium (*EDNRB1*), and pericyte marker (NG2).

#### Figure S2

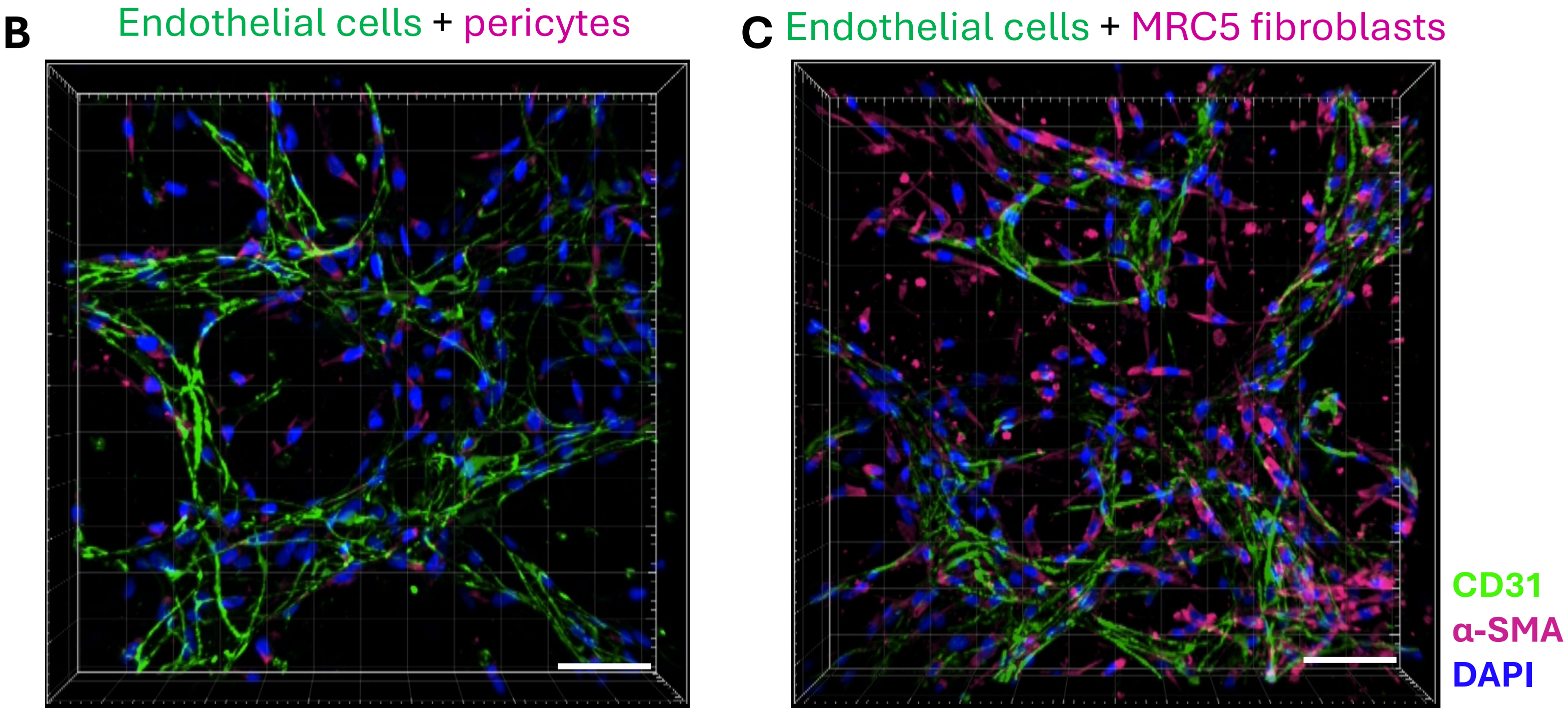


**Figure S2. Comparison of endothelial network formation with pericytes and fibroblasts.** Representative confocal image after 6 days of endothelial vascular network formation with pericytes **(B)** and MRC5 fibroblasts **(C)** stained for endothelial cell marker CD31 (in green) and in α-SMA (in magenta), marker for both pericyte and fibroblasts and nuclei with DAPI stain (in blue). Scale Bar, 20 μm.

#### Supplementary Figure 3

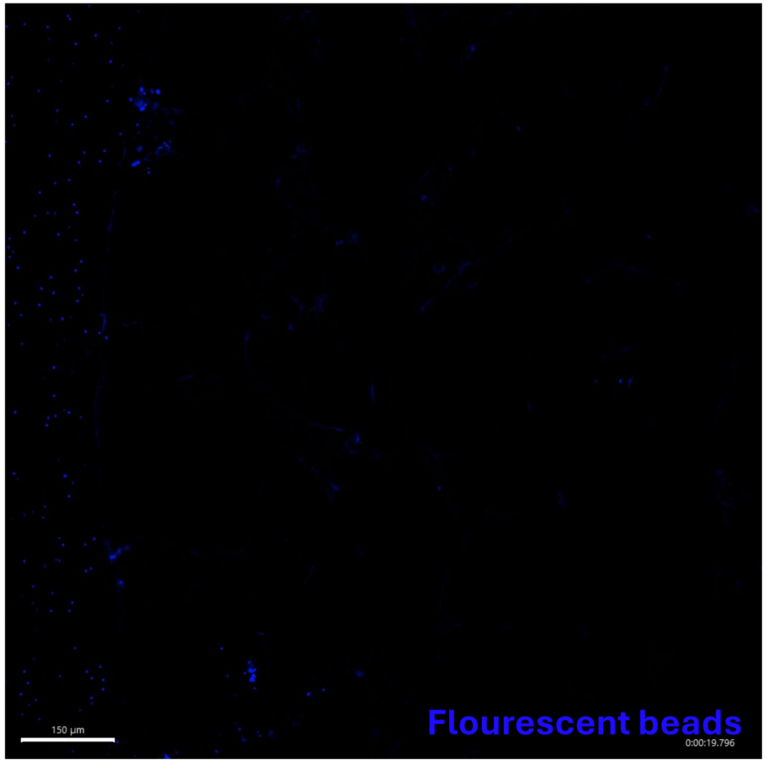


**Supplementary Figure 3.** Representative fluorescent image of a non-perfusable vascular network, characterized by the accumulation of 2 µm fluorescent beads (blue) at the pillar in the inlet, indicating obstruction and lack of bead entry into the network. Scale bar, 150 µm
