## Supplementary Figure legends and materials for "Modelling cigarette smoke-induced lung vascular dysfunction using an alveolus-on-chip"

**Table S1. List of primary and secondary antibodies used for staining**

| **S.No** | **Gene** | **Forward primer (5’-3’)** | **Reverse primer (5’-3’)** |
| --- | --- | --- | --- |
| 1 | *CD31* | AAATGCTCTCCCAGCCCAGGAT | GCAACACACTGGTATTCGACGTCTT |
| 2 | *LYVE1* | GGGTTGGAGATGGATTCGTGG | ATAGGCTGCAAACTGTCGGC |
| 3 | *CLDN5* | CTCTGCTGGTTCGCCAACAT | CAGCTCGTACTTCTGCGACA |
| 4 | *PRPRB* | TACCAATGGATCAACAGTGCC | GCATCAGCCGGTATCGTTCC |
| 5 | *GPIHBP1* | GCAACCTGACGCAGAACTG | CCAGGGTGGGACATTGCAC |
| 6 | *EDNRB1* | GTCCCAATATCTTGATCGCCAG | AAGGCACCAGCTTACACATCT |
| 7 | *ATP5B* | TCACCCAGGCTGGTTCAGA | AGTGGCCAGGGTAGGCTGAT |
| 8 | *RPL13A* | AAGGTGGTGGTCGTACGCTGTG | CGGGAAGGGTTGGTGTTCATCC |
| 9 | *NG2* | GCCACGTTGTCAGTCGATG | CCCATAGGGGACCTCTAGGG |

| **S.No** | **Antibody** | **Supplier** | **Catalog** | **Species** | **Dilution** |
| --- | --- | --- | --- | --- | --- |
| 1 | CD31 | Dako | #M0823 | mouse | 1:200 |
| 2 | Neuron-glial antigen 2 (NG2) | Abcam | #ab255811 | rabbit | 1:800 |
| 3 | pro-surfactant protein C | Milllipore | #ab3786 | rabbit | 1:100 |
| 4 | Cleaved caspase-3 | Cell signaling | #9664 | rabbit | 1:400 |
| 5 | Periaxin | Sigma Aldrich | #HPA001868 | rabbit | 1:100 |
| 6 | AF488 donkey anti-mouse | Thermo Fischer Scientific | #A-21202 | donkey | 1:400 |
| 7 | AF647 donkey anti-rabbit | Thermo Fischer Scientific | #A-31573 | donkey | 1:400 |
| 8 | Mouse IgG1 isotype control | R&D systems | #MAB002 | mouse | 1:200 |
| 9 | Rabbit IgG isotype control | Biotechne | #NBP2-24891 | rabbit | 1:100 |

**Table S2. List of forward and reverse primers used for qPCR analysis**

**Supplementary figures:**

**Fig S1. (A)** Flow cytometry analysis of endothelial cells isolated from lung tissue and expanded at P1, for CD140b/PDGRFβ and CD31 (n=3 different donors). **(B)** Bright field images (of n=3 different donors) depicting morphology of endothelial monolayers expanded at P1. **(C)** Vascular network formation of endothelial cells isolated from COPD donor. Representative confocal image (n=2) after 6 days of self assembled vascular network stained with endothelial cell marker, CD31 (in green) and pericyte marker NG2 (in red) and nuclei with DAPI stain (in blue). Scale bar 70 µm. (**D**) Flow cytometry analysis of endothelial cells inefficiently isolated with mesenchymal contamination, from lung tissue and expanded at P1, for CD140b/PDGRFβ and CD31 (n=1). (**E**) Bright field images (of n=1) depicting morphology of endothelial monolayers in the presence of mesenchymal cells, expanded at P1. **(F)** Representative confocal image after 6 days of self-assembled vascular network (n=3 different donors) stained with pericyte marker NG2 (in red) and nuclei with DAPI stain. Scale bar 70 µm.

**Fig S2. Gene expression analysis of pericyte monolayer. (A)** qPCR analysis of mRNA expression in pericytes (*n*=1) and endothelial cells (*n*=1) for markers of endothelial cells (*CD31*), macrovascular endothelium (*LYVE1*), microvascular endothelium (*CLDN5*), gCAP-microvascular endothelium (*GPIHBP1*, *PTPRB*), alveolar repair marker (*BMP6*), aCAP-microvascular endothelium (*EDNRB1*), and pericyte marker (NG2).

**Fig S3.** Representative fluorescent image of a non-perfusable vascular network characterised by the accumulation of 2µm fluorescent beads (blue) at the pillar of the inlet, indicating obstruction and lack of bead entry into the network. Scale bar 150 μm

**Supplementary video legends:**

**Video S1.** Representative confocal video (n=3 different donors) after 6 days of a self-assembled vascular network formation on chip with endothelial cells in green (CD31), pericytes in red (NG2), and nuclei in blue (DAPI). Scale Bar, 50 µm.

**Video S2.** Representative fluorescent video of self-assembled vascular network perfused using fluorescent beads (blue, 2 µm diameter). Scale bar, 100 µm.

**Video S3.** Representative fluorescent video of self-assembled vascular network (red - phalloidin) perfused using fluorescent beads (blue, 2 µm diameter) and live THP-1 monocytes (green). Scale bar, 150 µm.

**Video S4.** Representative confocal video (n=3 different donors) of alveolus on-chip with self-assembled vascular network stained for CD31 (in green) and AEC2 monolayer stained for (pro-SFTPC) (in red) and nuclei stained with DAPI (in blue). Scale bar: 20 µm.
